# Transcriptome analysis of the response of domesticated emmer wheat (*Triticum turgidum* subsp. *dicoccum*) to single vs. mixed infections with *Zymosptoria tritici* and *Puccinia graminis*

**DOI:** 10.1101/2025.10.13.682012

**Authors:** Miguel Ángel Corrales Gutiérrez, Francesca Desiderio, Dimitar Douchkov, Salim Bourras

**Author notes:** **Corresponding author** Salim Bourras.

## Abstract

Tetraploid emmer *Triticum turgidum* subsp. *dicoccum* is one of the first wheat species to be domesticated. Emmer shares many pathogen species with modern durum wheat (*Triticum turgidum* subsp. *durum*) and hexaploid bread wheat (*Triticum aestivum* L.). Domesticated emmer is considered a highly valuable source of potentially novel disease resistance genes that can be introduced into modern wheat varieties, however, large gaps remain in our understanding of the basic immune responses of emmer to the most common wheat pathogens. In this work we describe the transcriptional response of emmer to single and mixed pathogen species infections with the ascomycete *Zymoseptoria tritici*, and the basidiomycete *Puccinia graminis*. In doing so, we here uncover unique patterns in the response of emmer to a more complex pathogen population with mixed species infections.

## Introduction

Tetraploid emmer wheat (*Triticum turgidum* subsp. *dicoccum*) represents one of the earliest domesticated wheat species and served as a foundational crop during the agricultural revolution in the Fertile Crescent. Although emmer has been largely replaced by modern durum and bread wheat varieties in contemporary agriculture, it remains an invaluable genetic resource for crop improvement (Avni et al. 2017). Domesticated emmer shares many pathogen species with its modern relatives and is considered to harbor potentially novel disease resistance genes that could be introduced into elite wheat cultivars through breeding programs (Liu et al. 2017; Peleg et al. 2009). However, despite its recognized value as a source of resistance traits, significant gaps remain in our fundamental understanding of how emmer wheat responds to common wheat pathogens at the molecular level.

Two of the most economically important fungal diseases affecting wheat worldwide are septoria tritici blotch and stem rust, caused by *Zymoseptoria tritici* (*Z. tritici*) and *Puccinia graminis* f. sp. *tritici* (*P. graminis*) respectively. These pathogens employ markedly different infection strategies and occupy distinct ecological niches within the host plant. *Zymoseptoria tritici* is a hemibiotrophic pathogen that colonizes the apoplastic space systemically throughout leaf tissue, initially growing as an endophyte before transitioning to destructive necrotrophic growth (Rudd et al. 2015; Fones and Gurr 2015). In Europe alone, septoria tritici blotch represents the most economically damaging foliar disease of wheat, with yield losses reaching up to 50% in susceptible cultivars and accounting for approximately 70% of fungicide expenditure in the European Union (Fones and Gurr 2015). In contrast, *Puccinia graminis* f. sp. *tritici* is an obligate biotrophic pathogen that forms specialized feeding structures called haustoria, primarily in epidermal cells, and remains largely restricted to the leaf surface and immediate underlying tissue (Duplessis et al. 2011). These fundamental differences in pathogen biology suggest that host plants may deploy distinct defense strategies against each pathogen, yet how these responses are orchestrated at the transcriptional level in emmer wheat remains poorly characterized.

In natural agricultural settings, wheat plants are frequently exposed to multiple pathogens simultaneously or in rapid succession, leading to complex mixed infections. Understanding how plants respond to such mixed pathogen populations is critical for developing durable disease resistance strategies, yet most molecular studies of plant-pathogen interactions have focused exclusively on single pathogen infections under controlled conditions. The transcriptional response of wheat to mixed infections may differ substantially from responses to individual pathogens, potentially revealing emergent patterns in immune responses or unexpected interactions between different defense pathways. Furthermore, when pathogens with contrasting infection strategies co-occur, it remains unclear whether the host response represents a simple additive effect, whether one pathogen response dominates, or whether entirely novel response patterns emerge.

In this study, we characterized the transcriptional response of domesticated emmer wheat to single and mixed infections with *Z. tritici* and *P. graminis* using RNA sequencing. Our objective was to identify patterns of gene expression associated with each infection type and to determine how the host transcriptional response to mixed infections relates to responses elicited by each pathogen individually. Through comparative transcriptomic analysis, we provide insights into the complexity of emmer wheat immune responses under conditions that more closely approximate the multi-pathogen environments encountered in agricultural production systems.

## Methods and Materials

### Experimental Design

The experimental design aimed to analyze host-pathogen interactions during single and simultaneous infections with multiple pathogens. Twenty 9-day-old wheat seedlings were inoculated with *Z. tritici* at a concentration of 1×10^6^ spores/ml, while an additional ten plants of the same age were inoculated with 0.01% Tween-20 as a mock control. At three days post-inoculation with *Z. tritici*, both groups were subdivided into two cohorts of ten plants each. One cohort from each group was subsequently inoculated with *P. graminis* urediniospores at 250 spores/μl, while the other cohort received a second mock inoculation with 0.01% Tween-20. This design yielded four treatment groups: mock control, single infection with *Z. tritici*, single infection with *P. graminis*, and mixed infection with both pathogens.

### Plant Material and Growth Conditions

Seeds of the domesticated emmer wheat cultivar MG5323 (*Triticum turgidum* subsp. *dicoccum*) were obtained from Dr. Francesca Desiderio at the Council for Agricultural Research and Economics (CREA), Italy. Seeds were germinated in pots containing sterilized substrate and maintained under controlled environmental conditions at 20-21°C with 69-70% relative humidity. Plants were grown in a phytotron chamber under a photoperiod of 18 hours light and 6 hours darkness with a light intensity of 300 μmol m^−2^ s^−1^.

### Fungal Material and Inoculum Preparation

#### *P. graminis* Inoculum

*P. graminis* were collected from naturally infected wheat fields in Sweden during the summer of 2021. The isolate was genotyped using universal ITS primers (White et al. 1990) and propagated on the susceptible wheat cultivar Morocco. Spores were stored under dry conditions at 4°C until use. For inoculation, 6 mg of urediniospores were suspended in 1 ml of 0.01% Tween-20, and spore concentration was confirmed by hemocytometer counting to achieve a final concentration of 250 spores/μl.

#### *Z. tritici* Inoculum

Seven *Z. tritici* isolates (ST99CH_3B8, ST99CH_3C4, ST99CH_3C7, ST99CH_3D5, ST99CH_3D7, ST99CH_3F2, and ST99CH_3G6) originally collected in Switzerland in 1999 were obtained courtesy of Prof. Bruce McDonald at ETH Zürich. Isolates were cultured on yeast extract-malt extract-starch agar medium (YMS: 4 g/L yeast extract, 4 g/L malt extract, 4 g/L sucrose, 16 g/L agar) and incubated in darkness at 20°C for 11 days. Spores were harvested by scraping cells from plates under sterile conditions and resuspending them in 0.01% Tween-20. The suspension was filtered through a sterile filter with a pore diameter of 22-25 μm. Spore concentration was determined using a hemocytometer and adjusted to 1×10^6^ spores/ml for each isolate. Prior to inoculation, all seven isolates were mixed in equal volumes to create a composite inoculum.

### Pathogen Inoculation Protocol

#### *Z. tritici* Inoculation

Nine-day-old wheat seedlings were placed in a closed humidity chamber. The composite *Z. tritici* inoculum was applied using a spray gun at a rate of 0.33 ml per seedling. Mock control plants were inoculated with 0.01% Tween-20 using the same application method. Following inoculation, plants were maintained in the humidity chamber at greater than 70% relative humidity in complete darkness for 24 hours. Plants were then returned to the standard day-night cycle described above and maintained at high relative humidity through daily misting with distilled water using a low-pressure spray gun.

#### *Puccinia graminis* Inoculation

At three days post-inoculation with *Z. tritici* or mock treatment, plants designated for *P. graminis* infection were inoculated by applying the urediniospore suspension (250 spores/μl) directly to all leaf surfaces using sterile brushes until complete coverage was achieved. Control plants received 0.01% Tween-20 applied in the same manner. All plants were maintained under the standard photoperiod and high relative humidity conditions until sampling at 12 days post-inoculation with *Z. tritici*.

### RNA extractions and sequencing

#### RNA Extraction and Quality Assessment

Leaf tissue samples were collected at 12 days post-inoculation with *Z. tritici* (9 days post-inoculation with *P. graminis* for coinfected samples). Total RNA was extracted using the miRNeasy kit (Qiagen, Hilden, Germany) according to the manufacturer’s protocol. RNA quality was assessed by agarose gel electrophoresis to verify RNA integrity and using a NanoDrop ND-1000 spectrophotometer (Thermo Fisher Scientific, Waltham, MA, USA) to determine concentration and purity based on 260/280 nm absorbance ratios. RNA integrity was further evaluated using an Agilent Bioanalyzer 2100 (Agilent Technologies, Santa Clara, CA, USA).

#### RNA Sequencing

Total RNA (1–2 µg per sample) was used as input for library construction with the TruSeq Stranded mRNA Library Preparation Kit (Illumina, San Diego, CA, USA; RS-122-2101) according to the manufacturer’s protocol. Briefly, polyadenylated RNA was purified using oligo(dT) magnetic beads, followed by fragmentation at 94 °C for 8 minutes in first-strand synthesis buffer. First-strand cDNA synthesis was performed using SuperScript II Reverse Transcriptase and random hexamer primers at 25 °C for 10 minutes, 42 °C for 50 minutes, and 70 °C for 15 minutes. Second-strand synthesis was conducted with DNA Polymerase I and RNase H at 16 °C for 1 hour, incorporating dUTP in place of dTTP to enable strand specificity. Double-stranded cDNA was end-repaired, A-tailed, and ligated to TruSeq LT indexed adapters. Libraries were enriched by 12 cycles of PCR using Phusion High-Fidelity DNA Polymerase with the following cycling conditions: 98 °C for 30 s; 12 cycles of 98 °C for 10 s, 60 °C for 30 s, 72 °C for 30 s; and a final extension at 72 °C for 5 minutes. Samples were sequenced on NovaSeqXPlus (control-software 1.1.0.18335) at the SciLifeLab facility in Stockholm, Sweden, with a 151nt(Read1)-19nt(Index1)-10nt(Index2)-151nt(Read2) setup using ‘10B’ mode flowcell. The Bcl to FastQ conversion was performed using bcl2fastq_v2.20.0.422 from the CASAVA software suite. The quality scale used is Sanger / phred33 / Illumina 1.8+. The output generated ranged between 26 and 32 million reads (Mreads), with an average of ca. 29 Mreads per library. Raw sequence data have been deposited in Zenodo under accession number 10.5281/zenodo.17337984.

### Bioinformatics and Statistical Analysis

#### Quality Control and Read Processing

Raw sequencing data quality was assessed using FastQC v0.11.9 (Andrews 2010) to evaluate read quality metrics, sequence duplication levels, and potential adapter contamination. Adapter sequences and low-quality bases were removed using Trimmomatic v0.39 (Bolger et al. 2014) with the following parameters: ILLUMINACLIP:TruSeq3-PE.fa:2:30:10 SLIDINGWINDOW:4:15 MINLEN:36. The ILLUMINACLIP function removed TruSeq adapter sequences with seed mismatches tolerance of 2, palindrome clip threshold of 30, and simple clip threshold of 10. The SLIDINGWINDOW function performed sliding window trimming with window size of 4 bases and required quality of 15, while MINLEN dropped reads shorter than 36 bases after trimming. Read processing and quality control procedures were performed following previously established protocols (Praz et al. 2018; Poretti et al. 2020).

#### Transcript Quantification

Transcript abundance was quantified using Salmon v1.4.0 (Patro et al. 2017) in quasi-mapping mode with the following key parameters: --validateMappings --gcBias --seqBias. The *Triticum turgidum* subsp. *durum* cultivar Svevo transcriptome was used as a reference GenBank assembly GCA_900231445.1). Salmon indexing was performed using the salmon index command with default k-mer length (k=31). Quantification was executed using the salmon quant command in paired-end mode (-l A for automatic library type detection), which employed Salmon’s dual-phase parallel inference algorithm with selective alignment validation (--validateMappings) to improve mapping accuracy. Bias correction models were applied for fragment GC content (--gcBias) and positional sequence biases (--seqBias). Salmon generates transcript per million (TPM) values and estimated counts for all transcripts across all libraries using its expectation-maximization algorithm. Raw count data have been deposited in Zenodo under accession number 10.5281/zenodo.17337984.

#### Differential Gene Expression Analysis

Differential gene expression analysis was conducted using the Bioconductor package edgeR v3.34.1 (Robinson et al. 2010) in R v4.1.0 following established pipelines for RNA-seq data analysis (Praz et al. 2018). Raw count data obtained from Salmon quantification were imported into edgeR using the tximport package to aggregate transcript-level estimates to gene-level counts. A DGEList object was created using the DGEList() function, incorporating count data and sample group information. Low-expression genes were filtered to retain genes with counts per million (CPM) > 10 in at least three of the nine libraries, using the filterByExpr() function with default parameters to retain genes with adequate expression levels across samples. Normalization for RNA composition was performed using the trimmed mean of M-values (TMM) method implemented through the calcNormFactors() function (Robinson and Oshlack 2010). This normalization approach accounts for differences in library composition and sequencing depth between samples. Differential expression testing employed a generalized linear model (GLM) framework. Common, trended, and tagwise dispersions were estimated using the functions estimate GLMCommonDisp(), estimateGLMTrendedDisp(), and estimateGLMTagwiseDisp() respectively. The negative binomial generalized log-linear model was fitted to read counts for each gene using the glmFit() function, and differential expression was tested using likelihood ratio tests with the glmLRT() function through pairwise comparisons between treatment groups and the mock control. Multiple testing correction was applied using the Benjamini-Hochberg method to control the false discovery rate (FDR). Genes were considered differentially expressed if they exhibited a log_2_ fold change |log_2_FC| ≥ 2 and an adjusted p-value (FDR) ≤ 0.01. The topTags() function was used to extract ranked lists of differentially expressed genes for each comparison.

#### Gene Ontology Enrichment Analysis

Gene ontology (GO) enrichment analysis was performed using the topGO package in R (Alexa and Rahnenfuhrer 2021) to identify functional categories overrepresented among differentially expressed genes. GO term annotations for the durum wheat transcriptome were obtained from the Triticeae Multi-omics Center database. Enrichment testing was conducted using Fisher’s exact test with the runTest() function, employing the “weight01” algorithm to account for GO term hierarchical structure. GO categories were considered significantly enriched at p-value < 0.01 after multiple testing correction. Gene set enrichment analysis (GSEA) was conducted using the topGO package with the ‘preranked’ option on log_2_ fold-change-ranked genes against Gene Ontology gene sets for *Triticum turgidum* subsp. *durum* (Subramanian et al. 2005).

#### Data Visualization and Statistical Analysis

Multidimensional scaling (MDS) analysis was performed using the plotMDS() function in edgeR to visualize overall patterns of gene expression similarity across samples and treatments. For visualization of highly expressed genes and global expression patterns, counts were converted to reads per kilobase million (RPKM) and displayed as heatmaps using the pheatmap R package. Volcano plots were generated using the EnhancedVolcano package (Blighe et al. 2018) to display the distribution of differentially expressed genes based on log_2_ fold change and statistical significance (-log10 adjusted p-value).

## Results

### Data description and multidimentional scaling

A total of 12 samples were applied to Illumina NovaSeqXPlus, 150 base pairs, paired-end sequencing, corresponding to three treatments and one uninfected control. Libraries ranged between 26 and 32 Mreads, with an average of 29 Mreads per replicate per sample. Transcripts were quantified using Salmon (Patro et al. 2017) and the *Triticum turgidum* subsp. *durum* cultivar Svevo transcriptome as a reference (Maccaferri et al., 2019; Vendramin et al. 2019) (see Methods). Based on this initial mapping, the ‘Reads Per Kilobase of transcript per Million mapped reads’ (RPKM) values, and ‘Counts Per Million’ (CPM) values were calculated for all transcripts from all libraries and used to conduct a multidimentional scaling (MDS) analysis. The resulting data showed that RPKM and CPM counts were consistent across all replicates and all samples (see ‘quant’ output files in supporting data). For the MDS analysis, while samples grouped differently depending on the input i.e. RPKM (Figure 1A) or CPM (Figure 1B), samples corresponding to single infection with *P. graminis* (P) consistently grouped together with the uninfected mock control (M). Conversely, samples corresponding to single infection with *Z. tritici* (Z), grouped together with those infected with both pathogens (ZP) (Figure 1A,B). This data suggests that on the one hand infection with *Z. tritici* results in much stronger transcriptional changes than *P. graminis* (P) as compared to the mock control (at least in our experimental conditions). Which, on the other hand, seems to mask the effect of *P. graminis* in the mixed infections (ZP) where the samples are transcriptionally more similar to single infection with *Z. tritici* (Z).

**Figure 1.**
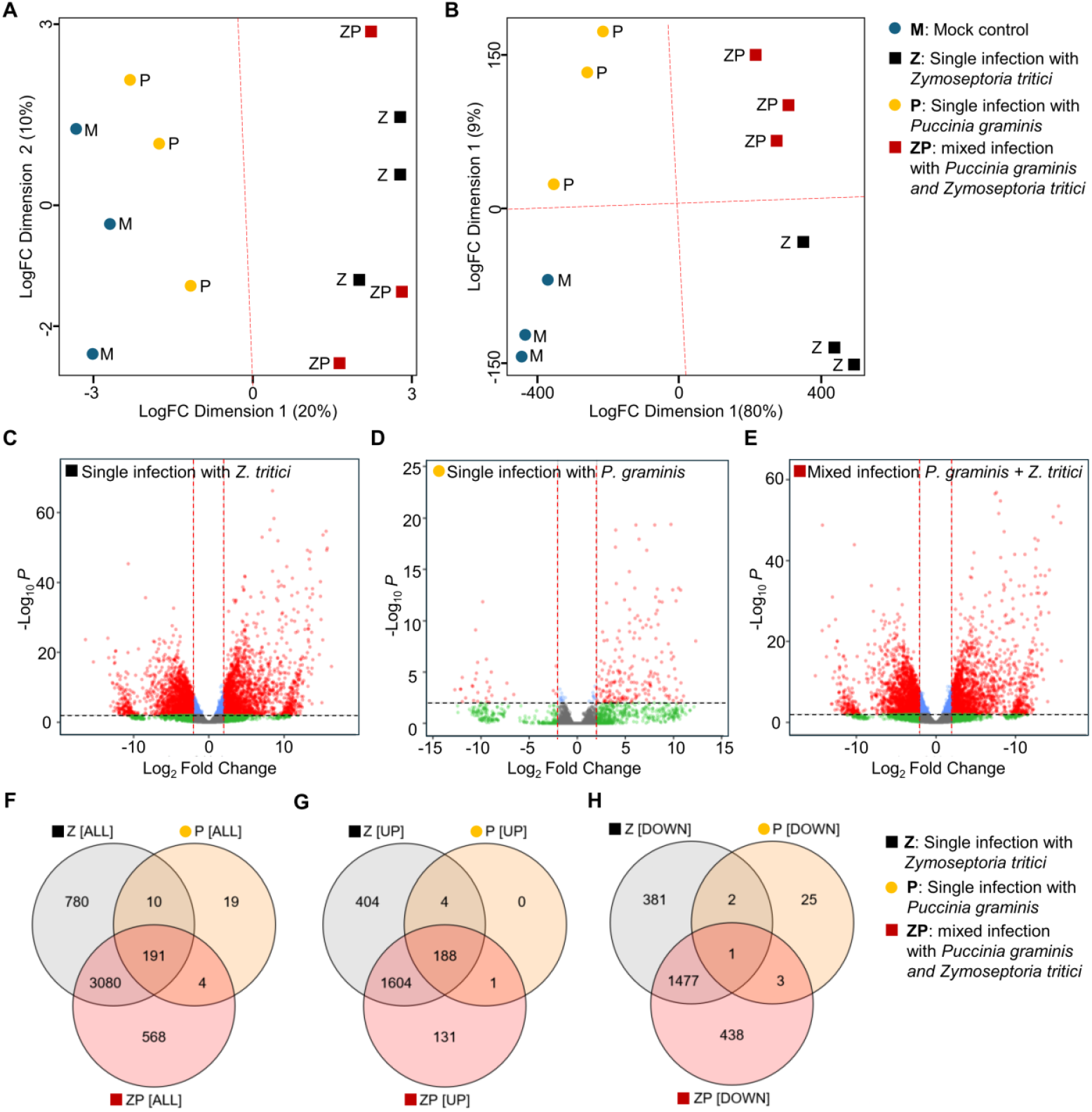
Multidimensional scaling (MDS) analysis and differential gene expression overview. (A) MDS plot based on RPKM values showing sample clustering by treatment. (B) MDS plot based on CPM values demonstrating consistent grouping patterns where *P. graminis* single infection (P) clusters with mock control (M), while *Z. tritici* single infection (Z) groups with mixed infection (ZP). (C-E) Volcano plots showing differentially expressed genes (log_2_ fold change ≥ 1, adjusted p-value ≤ 0.05) for each treatment compared to mock control: (C) *Z. tritici* single infection, (D) *P. graminis* single infection, and (E) mixed infection. Red dots indicate upregulated genes, blue dots indicate downregulated genes, and gray dots represent genes below significance thresholds. (F-H) Venn diagrams comparing differentially expressed genes between treatments: (F) all differentially expressed genes, (G) upregulated genes only, and (H) downregulated genes only. Numbers indicate unique and shared gene sets, demonstrating that mixed infection shares substantially more genes with *Z. tritici* single infection (3,080) than with *P. graminis* single infection (10).

### Differential gene expression analysis

Differential gene expression analysis was performed using the Bioconductor package edgeR (Robinson et al. 2010) using normalized RPKM counts as previously described (Praz et al., 2018; Poretti et al., 2020). Differentially expressed genes passing a Log_2_Fc threshold ≥ 1 and a p-value ≤ 0.05 in comparison to the uninfected control were visualized as a volcano plot (see Methods) (Figure 2C,E). Consistent with the MDS analysis, differences between the *P. graminis* treatment (Figure 2D) and both treatments implicating *Z. tritici* were striking (Figure 2A,E). The volcano plots further show that infection with *Z. tritici* resulted in much stronger transcriptional changes than the obligate *P. graminis*, to such an extent that mixed infections with both pathogens seem to be transactionally dominated by the host response to *Z. tritici*. Consistent with this, comparison of differentially expressed, upregualted, and downregulated genes between treatments showed that wheat response to mixed infection shared 3080 unique differentially expressed genes with the *Z. tritici* treatment, and only 10 with the *P. graminis* treatment (Figure 1F). The same pattern is observed when considering upregulated and downregulated genes separately (Figure 1G,H). These results further confirm that the emmer wheat response to mixed infections with *Z. tritici* and *P. graminis* is more similar to the response to single infection with *Z. tritici*, and arguably different from single infection with *P. graminis*.

**Figure 2.**
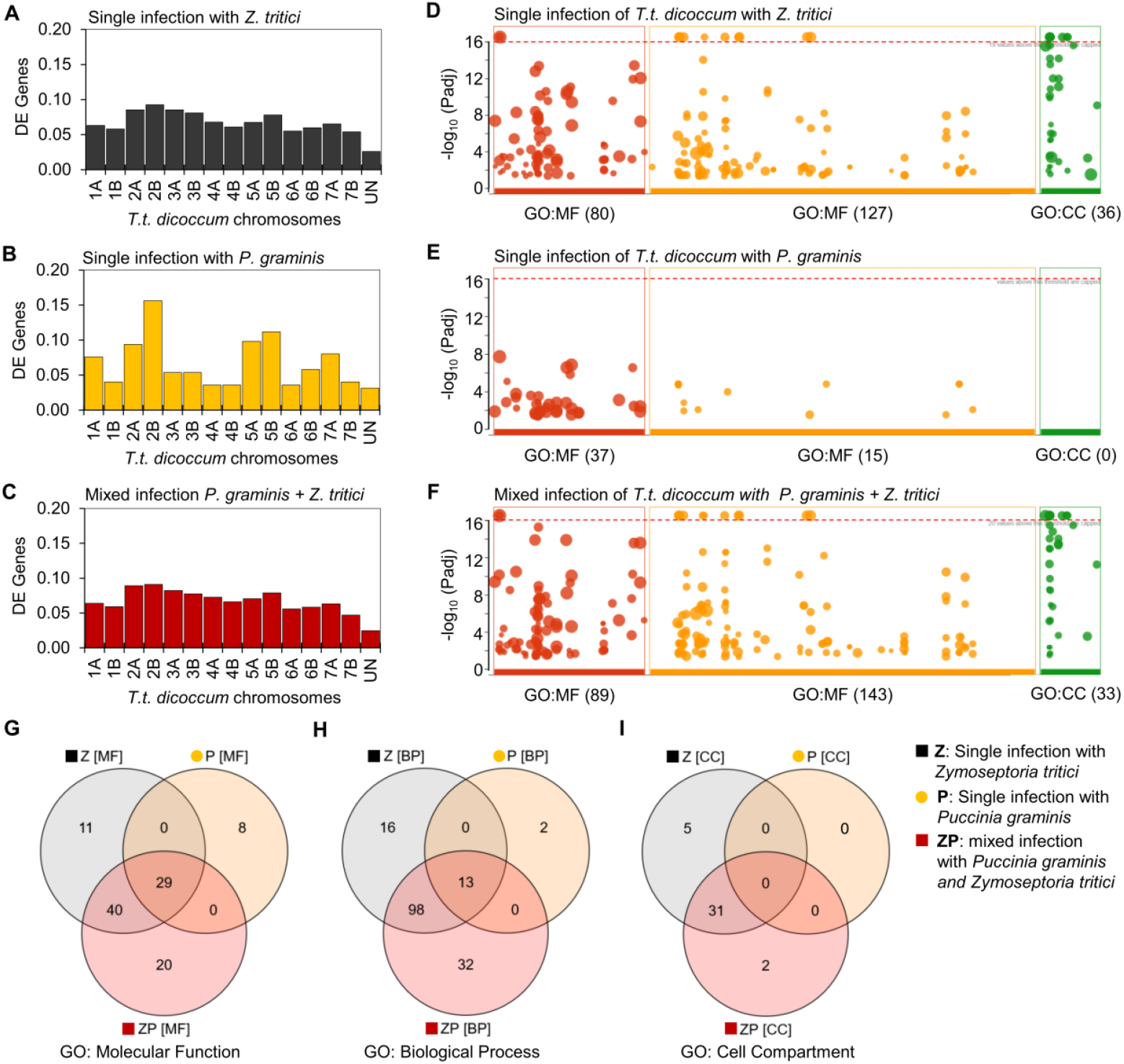
Chromosome-wide distribution of differentially expressed genes and gene ontology enrichment analysis. (A-C) Distribution of differentially expressed genes across all 14 chromosomes (1A-7A, 1B-7B) and unassembled contigs (UN) of the tetraploid emmer wheat genome for: (A) *Z. tritici* single infection, (B) *P. graminis* single infection, and (C) mixed infection. Note the distinct patterns on chromosomes 2A, 2B, 5A, and 5B in *P. graminis* treatment compared to *Z. tritici*-containing treatments. (D-F) Gene Ontology enrichment analysis showing significantly enriched terms (p-value < 0.01) in three categories: Molecular Function (MF), Biological Process (BP), and Cellular Component (CC) for: (D) *Z. tritici* single infection, (E) *P. graminis* single infection (showing no significant enrichment), and (F) mixed infection. (G-I) Venn diagrams comparing GO term overlaps between treatments for: (G) Molecular Function, (H) Biological Process, and (I) Cellular Component categories, revealing both shared and treatment-specific functional annotations.

We suggest that such differences are possibly reflecting differences between the invasive strategies (at the time of sample collection) between these two pathogens where **(i)** *Z. tritici* is a hemibiotroph and a systemic apoplastic colonizers of the whole leaf tissue, while **(ii)** *P. graminis* is an obligate biotroph mostly restricted to epidermal cells through the formation of feeding structure (haustoria) and mostly occupying the epiphytic space. We also acknowledge that this could also be partially explained by differences in inoculum load and inoculation success under controlled lab conditions, which would also highlight a possible limitation of such a complex artificial infection scheme.

### Chromosome-wide distribution of differentially expressed genes

We then compared the distribution of differentially expressed genes between treatments across all chromosomes of the tetraploid durum wheat genome (Chromosomes 1A-7A, 1B-7B) as well as the unassembled contigs (UN) (Figure 2A-C). While the distribution of differentially expressed genes was very similar between single infections with *Z. tritici* (Figure 2A), and samples with mixed infections (Figure 2C), the difference with single infection *P. graminis* was striking (Figure 2B). Chromosomes 2(A,B) and 5(A,B) in particular, seem to have a much higher contribution to the emmer wheat response to *P. graminis* alone than in the two other treatments with *Z. tritici*. Overall, the low number of differentially expressed genes in the *P. graminis* treatments can be partially explained by the biotrophic lifestyle of the pathogens (see previous section). Therefore, we suggest that the host response to *P. graminis* in this assay must be functionally different from the host response to mixed infection, which is likely masked by the overwhelming response to *Z. tritici*.

### Enrichment analysis

To gain functional insight into the differences between treatments, we applied differentially expressed genes to functional annotation using the Gene Ontology classification (Ashburner et al., 2000) (Supplementary Table R2), followed by a gene set enrichment analysis (GESA) (Figure 2D-F) (Subramanian et al., 2005). We found no enrichment in any of the GO categories for the treatment with *P. graminis* alone, while the category Cell Compartment (CC) had no annotations (Figure 2E). This could be explained by the lifestyle of the pathogen and the small number of differentially expressed genes found in the first place. For the treatment with mixed infections and *Z. tritici* alone, a clear functional signature was revealed with significant gene set enrichments in all three GO categories (Figure 2D,F). Further comparisons of the GO terms between all treatment (Figure 2 G-I) show that the functional response of emmer wheat to *P. graminis* is largely encompassed within a core response of to all treatments (i.e. central intersection in the Venn diagram) (Figure 2G,H). In contrast, and despite large similarities shown in previous analyses, treatments with *Z. tritici* alone and in mixed infection with *P. graminis* had both shared and treatment specific GO annotations, in all categories (Figure 2 G-I). This data suggests that there is a common response of emmer wheat to all treatments. However, while the presence of *Z. tritici* seems to mask the host response to *P. graminis* in mixed infections, these two responses are not identical and could nevertheless still be distinguished at the functional level.

### Comparison of candidate pathways

Finally, we analyzed the GO data for candidate pathways that could provide additional insights into the host response based on the most significantly enriched gene sets in the category Biological Function (BP) (Figure 3 A-D). The analysis could therefore only include the treatments with *Z. tritici* alone and in coinfection with *P. graminis* (Figure 2D,F), considering that no enrichment in any GO category was found for the treatment with *P. graminis* alone (Figure 2E). We found that the most significant gene set enrichments were in hallmarks of plant defense to pathogens, with the most prominent examples being the carboxillic acid metabolism pathway, and the oxoacid metabolism pathway, which are enriched in the response to *Z. tritici* alone and in combination with *P. graminis*. Closer analysis of gene expression heatmaps show distinct gene activation patterns clearly differentiating the host response to the treatments where

**Figure 3.**
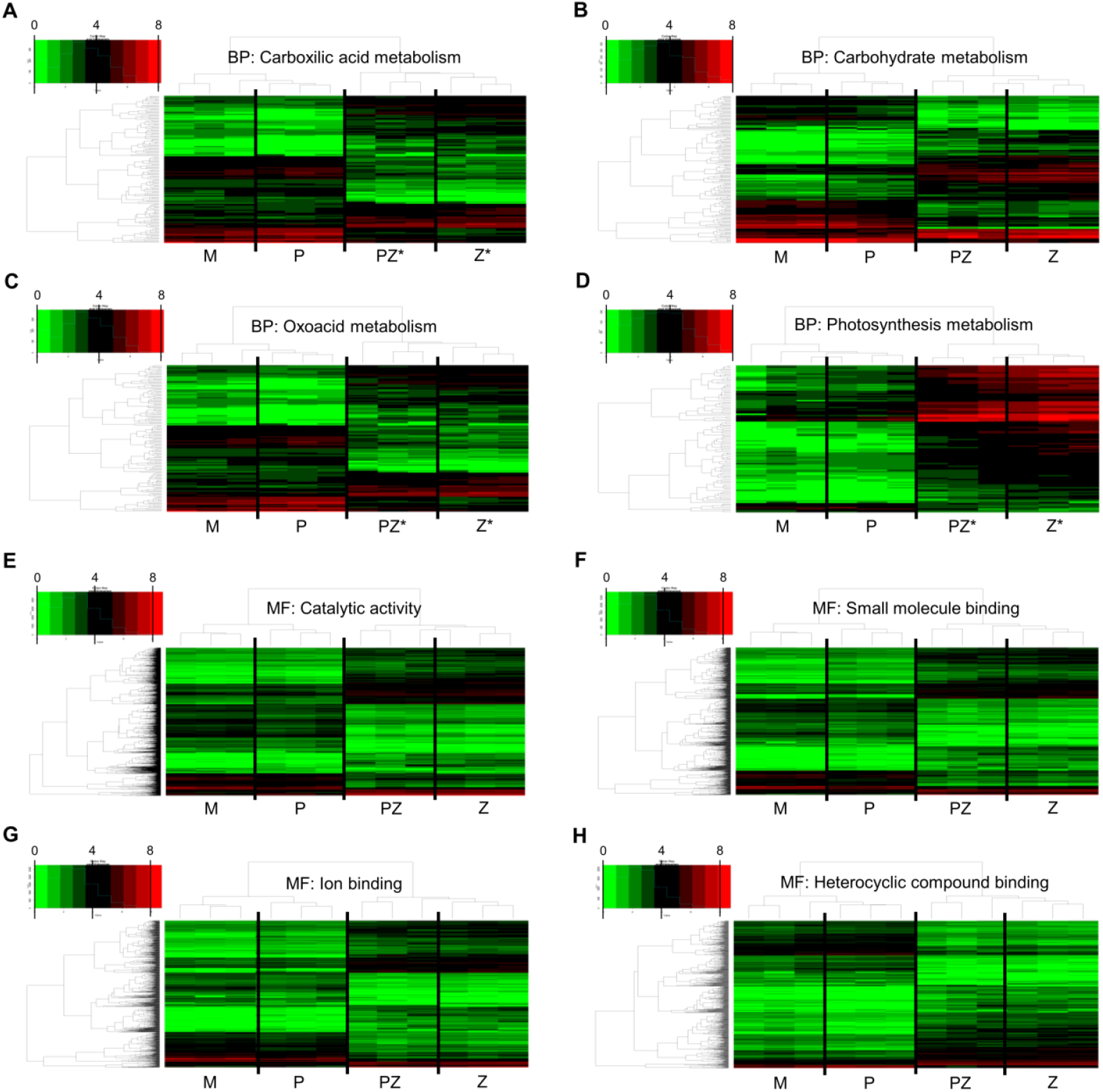
Candidate pathway analysis of key biological processes. (A-D) Heatmaps showing expression patterns of genes associated with the most significantly enriched biological processes from GO analysis: (A) carboxylic acid metabolic process, (B) oxoacid metabolic process, (C) carbohydrate metabolic process, and (D) photosynthesis. Each heatmap displays normalized gene expression values across treatments (Mock, *P. graminis, Z. tritici*, Mixed infection) with genes clustered by expression similarity. Color scale ranges from green (low expression) to red (high expression). These pathways represent hallmarks of plant defense responses and metabolic reprogramming, showing clear patterns that distinguish treatments containing *Z. tritici* from *P. graminis* alone or mock control. The carboxylic acid and oxoacid metabolism pathways are particularly relevant for antimicrobial compound biosynthesis and defense signaling, while alterations in carbohydrate metabolism and photosynthesis reflect the systemic metabolic changes associated with *Z. tritici* infection during the biotrophic-to-necrotrophic transition phase.

*Z. tritici* is present from *P. graminis* alone or the control (Figure 3 A,B). The carboxillic acid is prominent player of plant defensese against pathogens, where it can act as the precursors of active antimicrobial compounds, a defense signaling molecule, an elicitor of plant immune responses, or act directly as a toxin against pathogens (Morgunov et al. 2017; Godlewska-Żyłkiewicz et al. 2020; Seybold et al. 2020). Similarly, derivatives of the oxoacid metabolism are also hallmarks of plant defenses against pathogens, with the prominent example of oxylipins such as Jasmonic Acid (JA) (Eckardt 2008; Dave and Graham 2012; Liu and Park 2021). Another important derivative of the oxoacid metabolism is the 12-oxo-Phytodienoic acid which is proposed to play an important role in regulating the tradeoff between plant growth and plant defense (Liu and Park 2021).

Two additional terms stood out in the GESA with clear activation patterns differentiating the host response to the treatments where *Z. tritici* is present from *P. graminis* alone or the control: ‘Carbohydrate metabolism’ and ‘Photosynthesis’ (Figure 3 C,D). In fact, both pathways were shown to be significantly altered by *Z. tritici* infection, suggesting they are key targets for successful host colonization (Rudd et al. 2015; Palma-Guerrero et al. 2016). Furthermore, both pathways were shown to be particularly important for the switch from to the necrotrophic phase (Palma-Guerrero et al. 2016), thus coinciding with the sampling time point. In fact, it was shown that *Z. tritici* infection induces long-lasting systemic susceptibility and systemic reprogramming of the wheat metabolome (Seybold et al. 2020). Altogether, these results further substantiate the observation that infection with *Z. tritici* causes large systemic changes in the host that are likely to mask or alter infections with a secondary pathogen.

## Discussion

Our transcriptomic analysis reveals that *Z. tritici* dominates the transcriptional response in mixed infections with *P. graminis*, where co-infected emmer wheat plants closely resemble those infected solely with *Z. tritici*. This transcriptional dominance may be explained by the recently described systemic induced susceptibility (SIS), where *Z. tritici* can suppress host immune responses systemically, potentially creating conditions that affect secondary pathogen responses (Seybold et al. 2020). The strong *Z. tritici* transcriptional response likely reflects fundamental differences in infection strategies between the two pathogens. *Z. tritici* undergoes systemic apoplastic colonization and transitions from biotrophic to necrotrophic growth at our 12-day sampling timepoint (Rudd et al. 2015), coinciding with extensive host transcriptional reprogramming. Previous work showed that *Z. tritici* can suppress benzoxazinoids, phenylpropanoids, and lignin biosynthesis during this transition (Seybold et al. 2020). Our functional enrichment analysis reveals changes in carboxylic acid and oxoacid metabolism pathways, which are biochemically related to these defensive metabolites. In contrast, *P. graminis* maintains a more localized biotrophic interaction through haustorial feeding, potentially generating fewer systemic transcriptional changes.

The chromosome-wide analysis shows that *P. graminis* single infection produces distinct expression patterns on chromosomes 2A/B and 5A/B, indicating that emmer wheat possesses pathogen-specific resistance mechanisms consistent with characterized stem rust resistance genes located on these chromosomes (Karelov et al. 2022). The limited *P. graminis* transcriptional signature in mixed infections could reflect either the masking effect of the dominant *Z. tritici* response or potential interference with rust-specific immune pathways. However, the precise mechanisms underlying this interaction require further investigation. Importantly, our analysis reveals that co-infection does not simply replicate *Z. tritici* single infection. The presence of treatment-specific functional annotations suggests that plants retain some capacity for differential responses to mixed infections, indicating that the interaction between these pathogens and the host is more complex than simple transcriptional masking. This observation aligns with recent studies demonstrating that multi-pathogen interactions can produce unexpected disease outcomes that differ from predictions based on single-pathogen infections (Bellah et al. 2023).

Our results highlight the value of emmer wheat as a genetic resource for understanding pathogen interactions in ancestral wheat species. The distinct chromosomal distribution patterns of differentially expressed genes suggest that emmer wheat maintains pathogen-specific resistance architectures that may be valuable for developing varieties with enhanced resilience to multiple pathogens, particularly given the ongoing agricultural challenges posed by both septoria tritici blotch and stem rust diseases globally (Chai et al. 2022). These findings contribute to our understanding of multi-pathogen interactions in cereal crops and suggest that the transcriptional response to co-infection can be dominated by one pathogen while retaining pathogen-specific signatures. The transcriptional patterns we observe in emmer wheat provide insights into how ancestral wheat species respond to mixed pathogen infections and may inform strategies for improving disease resistance in modern wheat varieties. Future research should explore the mechanisms underlying pathogen dominance in co-infections and investigate whether similar patterns occur across different pathogen combinations and wheat species.

## Data Availability Statement

Raw RNA-sequencing data have been deposited as FastQ files in Zenodo under accession number 10.5281/zenodo.17337984. Raw count data have been deposited as full ‘qunant’ output files in Zenodo under accession number 10.5281/zenodo.17337984.

## Acknowledgments

We thank Prof Bruce A. McDonald (Department of Environmental Systems Science, ETH Zurich) for help and guidance with the selection of *Z. tritici* isolates. We thank Prof Daniel Croll (Laboratory of Evolutionary Genetics, Institute of Biology, University of Neuchâtel) for guidance with genome annotation resources for *Z. tritici*. We thank Fredric Hedlund and Ayano Tanaka for their technical assistance with the growth facilities at the SLU BioCentrum.

## Conflict of Interest

The authors have no conflicts of interest to declare.

## Funder Information

This work was supported by funding from the Swedish Research Council for Sustainable Development FORMAS (grant number 2020–01007), and the Carl Trygger Foundation (grant number 21:1171). Open access funding provided by the Swedish University of Agricultural Sciences.

**Table 1.** Summary of top 10 enriched GO terms.

| Single infection<br>with *Z. tritici* | Single infection<br>with *P. graminis* | Mixed infection with<br>*Z. tritici* + *P. graminis* |
| --- | --- | --- |
| **TOP 10 GO: Molecular Function** |  |  |
| Catalytic activity | Catalytic activity | Catalytic activity |
| Small molecule binding | Small molecule binding | Small molecule binding |
| Ion binding | Ion binding | Ion binding |
| Heterocyclic compound binding | Heterocyclic compound binding | Heterocyclic compound binding |
| Anion binding | Anion binding | Anion binding |
| Nucleotide binding | Carbohydrate derivative binding | Nucleotide binding |
| Nucleoside phosphate binding | Nucleotide binding | Nucleoside phosphate binding |
| Purine nucleotide binding | Nucleoside phosphate binding | Purine nucleotide binding |
| Carbohydrate derivative binding | Adenyl nucleotide binding | Carbohydrate derivative binding |
| Ribonucleotide binding | Purine nucleotide binding | Ribonucleotide binding |
| **TOP 10 GO: Biological Process** |  |  |
| Cellular process | Glucosamine-containing compound catabolic process | Cellular process |
| Metabolic process | Aminoglycan metabolic process | Metabolic process |
| Small molecule metabolic process | Aminoglycan catabolic process | Small molecule metabolic process |
| Carbohydrate metabolic process | Glucosamine-containing compound metabolic process | Carboxylic acid metabolic process |
| Carboxylic acid metabolic process | Amino sugar catabolic process | Oxoacid metabolic process |
| Oxoacid metabolic process | Chitin metabolic process | Organic acid metabolic process |
| Organic acid metabolic process | Chitin catabolic process | Carbohydrate metabolic process |
| Lipid metabolic process | Amino sugar metabolic process | Lipid metabolic process |
| Amino acid metabolic process | Cell wall macromolecule catabolic process | Amino acid metabolic process |
| Generation of precursor metabolites and energy | Cell wall macromolecule metabolic process | Generation of precursor metabolites and energy |
| **TOP 10 GO: Cell Compartment** |  |  |
| Cellular anatomical structure |  | Cytoplasm |
| Membrane |  | Plastid |
| Cytoplasm |  | Chloroplast |
| Plastid |  | Thylakoid |
| Chloroplast |  | Photosynthetic membrane |
| Thylakoid |  | Organelle envelope |
| Photosynthetic membrane |  | Plastid envelope |
| Organelle envelope |  | Thylakoid membrane |
| Plastid envelope |  | Plastid membrane |
| Thylakoid membrane |  | Bounding membrane of organelle |

## Notes

### Competing Interest Statement

The authors have declared no competing interest.

